# Are Antisense Proteins in Prokaryotes Functional?

**DOI:** 10.1101/2020.02.20.958058

**Authors:** Zachary Ardern, Klaus Neuhaus, Siegfried Scherer

## Abstract

Many prokaryotic RNAs are transcribed from loci outside of annotated protein coding genes. Across bacterial species hundreds of short open reading frames antisense to annotated genes show evidence of both transcription and translation, for instance in ribosome profiling data. Determining the functional fraction of these protein products awaits further research, including insights from studies of molecular interactions and detailed evolutionary analysis. There are multiple lines of evidence however that many of these newly discovered proteins are of use to the organism. Condition-specific phenotypes have been characterised for a few. These proteins should be added to genome annotations, and the methods for predicting them standardised. Evolutionary analysis of these typically young sequences also may provide important insights into gene evolution. This research should be prioritised for its exciting potential to uncover large numbers of novel proteins with extremely diverse potential practical uses, including applications in synthetic biology and responding to pathogens.

## Introduction

### The many functions of antisense RNAs

A wide range of non-coding RNAs have been characterised in bacterial genomes. Among these putatively non-coding sequences are many antisense transcripts. Indeed, up to 75% of all prokaryotic genes are associated with antisense RNAs - though the number differs significantly between species and according to the methods used (Georg and Hess, 2018). Their functions, if any, are poorly understood in most cases. The characteristics of antisense RNAs range widely in terms for instance of length, location in relation to the sense gene, and mechanisms of regulation (Lejars et al., 2019). In studies so far they are usually associated with reducing transcription of the sense gene, but they can also increase it, for instance by changing the structure of the sense transcript. They can influence single genes, or have global effects for instance through a target involved in general translation. Other known effects relate to functions including virulence, motility, various mechanisms of gene transfer, and biofilm formation (Lejars et al., 2019). The numerous examples of antisense transcription which have been investigated do not just include short antisense RNAs, though these are well-known; the many longer examples include a 1200 nucleotide antisense RNA in *Salmonella enterica*, AmgR (Courtney and Chatterjee, 2014). Antisense transcripts have been shown to be co-expressed within a single cell with the use of an antibody against double-stranded RNA in various studies, including in *Escherichia coli* and *Streptomyces coelicolor*, as reviewed in Georg & Hess (2018). Relatively little attention however has been paid to the possibility that RNA in antisense to protein coding genes may also frequently encode proteins (Georg and Hess, 2018). Rather than short, trivial overlaps, which are well known (Saha et al., 2016), here we focus on cases where an antisense (or “antiparallel”) ORF with evidence of translation is fully embedded within a known protein coding gene.

The existence of substantially overlapping gene pairs has been known since the beginning of modern genome sequencing, when the proteins directly detected in the bacteriophage phiX174 were shown to not be able to fit into the sequenced genome without the translation of overlapping open reading frames (ORFs) (Barrell et al., 1976). Since then, overlapping genes have typically been assumed to be fairly common only in viruses and extremely rare in other taxa, with the possibility of there being multiple examples in other taxa only sporadically discussed, e.g. Chou et al. (1996). However, their occurrence in bacteriophage in particular should raise the suspicion that they may be common in bacteria as well, given for instance the large amounts of genetic material transferred from temperate phage genomes to bacterial genomes (Harrison and Brockhurst, 2017, Owen et al., 2019). The properties of same-strand overlaps between viral genes have been studied, (Pavesi et al., 2018, Willis and Masel, 2018), but even in viruses, relatively little attention has been given to antisense overlaps. There is however increasing evidence for functional translated antisense open reading frames too, notably the antisense protein Asp in HIV-1 (Cassan et al., 2016, Affram et al., 2019, Nelson et al., 2019). In general it can be said that small ncRNAs are well recognized but their coding potential has generally been overlooked. Many might be protein-coding (i.e. mRNA), some are indeed ncRNA, and several will be dual-functional (e.g. (Neuhaus et al., 2017, Wadler and Vanderpool, 2007, Gimpel and Brantl, 2017). The same trichotomy of functional categories applies in the case of antisense RNAs.

In bacteria, a number of individual antisense proteins have been discovered; the lines of evidence for some of these will be discussed below. High throughput analyses of ribosome profiling data, which uncovers the part of the transcriptome associated with ribosomes (Ingolia et al., 2009), thus revealing the ‘translatome’, have begun to suggest that many more may be present. Friedman et al. (2017) found evidence for approximately 17 antisense ORFs, previously thought to be be non-coding sRNAs, translated over above the level expected by chance in *Escherichia coli* K12. The 10 sRNAs these belong to are shown in Figure 1A. Weaver et al. (2019) found ribosome profiling evidence, including evidence specifically for translation initiation (using retapamulin), for nine antisense overlapping gene candidates in *E. coli* K12, also shown in Figure 1A. As reported in a recent pre-print Smith et al. (2019) found many overlapping ORFs in *Mycobacterium tuberculosis* associated with ribosomes using retapamulin. From 355 novel ORFs expressed in two replicates they report 241 overlapping and embedded in annotated genes, including both sense and antisense overlaps, of which many were very short. Of those encoding at least 20 amino acids, 51 are antisense embedded. These antisense ORFs are shown in Figure 1B. However, it was claimed that as a class the novel ORFs are not under selection, and the association with ribosomes was attributed to non-functional pervasive translation (Smith et al., 2019). Jeong et al. (2016) reported ribosome profiling in *Streptomyces coelicolor* - although this result was not highlighted, examining the supplementary data showed 10 antisense putative sRNAs with ribosome profiling evidence. No doubt many more such discoveries await systematic analysis of published ribosome profiling data. There are also many putative same-strand overlapping genes, as discussed early on by e.g. (2003), showing that alternate frame translation is likely a general phenomenon - but these have also been claimed to not be under selection (Meydan et al., 2019). This increasing evidence for translation of both sense and antisense alternate frame ORFs, currently only typically acknowledged as ncRNAs, should push the question of ‘pervasive function’ and how to categorise the range of translated ORFs to the forefront of microbiology, but it is yet to receive substantial attention. The evidence of expression in alternate frames is generally ignored, and when acknowledged it is generally presumed to be non-functional - however, we argue this inference is made too quickly on insufficient grounds. Here we explore how to ascertain function and present a few examples of antisense genes with evidence for functionality.

**Figure 1.**
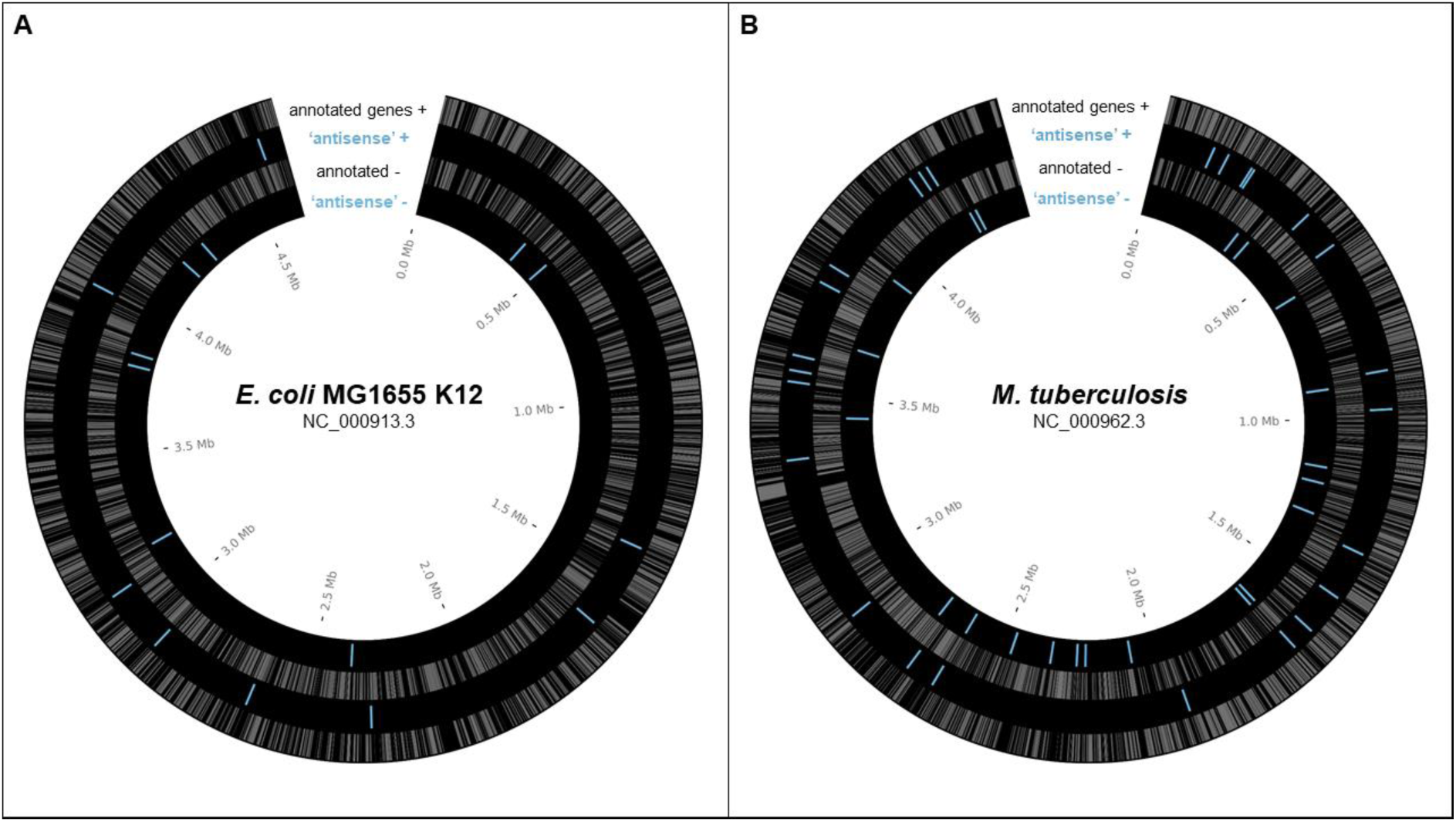
Reported potential protein-coding ORFs overlapping in antisense, based on ribosome profiling experiments. A) Reported antisense OLGs in E. coli K12 (NC_000913.3) - Weaver et al. 2019 & Friedman et al 2017 (sRNAs with evidence of translation). B) Reported antisense OLGs in M. tuberculosis (NC_000962.3) - Smith et al.2019. Annotated genes grey. Antisense blue.

### ‘Function’ in a biological context

The question of what counts as ‘function’ in a biological context is not straightforward. An interdisciplinary group of researchers have recently discussed the issue in relation specifically to *de novo* gene origin (Keeling et al., 2019), and proposed five categories of meanings of function, pertaining to expression, capacities, interactions, physiology and evolution. As they helpfully note, “Separating these meanings from one another enables communicating with increased precision about what the findings are, thereby helping to [avoid] fallacious logical shortcuts such as ‘this protein is expressed therefore it is functional therefore it is under selection’”. Interestingly, they had limited success in actually applying their categorisation, with most instances in a test set of article abstracts not uniformly assigned to a category by different team members. This suggests that biologists should write with more precision to clarify the sense of function intended. In this article we will focus on the senses relating to the biochemical “capacity” of the products of genetic elements and their evolutionary history, although the other senses will also come into play. The unifying general concept we use is that an element is functional if it does something useful for the organism in ecologically relevant circumstances.

The important philosophical questions have been reviewed elsewhere (Brandon, 2013). Here we summarise some established methods for determining function in the molecular biosciences and how they have been or could be applied to antisense proteins. It has become popular to adopt an etiological account of function i.e. that an element’s function depends on its selective history, particularly in relation to the dispute over how to assign function to elements in the human genome following the ENCODE project (Graur et al., 2013, Doolittle et al., 2014, Doolittle, 2018, Graur et al., 2015). However, the evolutionary etiology of biological systems is not always fully accessible to us (Ardern, 2018), and sometimes the history of selection in a lineage or for a particular gene may be inaccessible or the accessible parts incomplete in important ways. The genomic influence of different kinds of selection on bacterial genomes, including selective sweeps, background selection, positive selection, and purifying selection, remains a point of contention (Bendall et al., 2016, Takeuchi et al., 2015, Gibson and Eyre-Walker, 2019, Sela et al., 2019). Perhaps the most difficult issue here is how to characterize function in young genes, which may be somewhere along a spectrum between positive selection and purifying selection. Positive selection may be acting to modify a sequence which has only recently evolved or only recently become useful, for instance due to new environmental conditions. At some point however, modifications are overwhelmingly selected against, i.e. purifying selection is said to dominate. This fascinating transition region to our knowledge has received very little study, but it is plausible that most young genes fall within it and so are likely to be missed by methods seeking clear signatures of either purifying or positive selection. A recent study has shown that embedded overlapping genes in viruses usually evolve faster than the gene they are embedded in (Pavesi, 2019) - such cases will be missed by tests of purifying selection.

Additional relevant complexities include recombination, horizontal gene transfer, varying evolutionary rates, and unknown past environmental conditions. Evolutionary analyses certainly can provide strong evidence for function in cases of strong selection, but appropriate lower thresholds for determining that an element is functional while minimising false negatives are much harder to determine. Arguably of much greater relevance than etiology for molecular biologists is what a genetic element does in the current system, and whether it contributes to the goals or life-conducive activities of that system. That is, as the etiological theorists correctly emphasise, function is not just about ‘causal role’, it concerns a contribution to a wider system which is in some sense goal-directed. However given complex histories of multiple evolutionary forces this does not necessarily imply anything directly about a particular canonical signature of natural selection being observable in the existing sequence. A good example of these complexities is the prevalence of translation and likely functions in putative ‘pseudogenes’ (Goodhead and Darby, 2015, Cheetham et al., 2019).

### High-throughput experimental evidence

The ‘gold-standard’ proof of the active translation of a gene has traditionally been direct evidence from proteomics experiments, a technology which precedes modern genome sequencing by a few years. However, evidence from current proteomics methods is inherently limited even after decades of improvements. For instance, small proteins are notoriously difficult to detect by mass spectrometry, because upon proteolytic digestion they tend to generate no suitable peptides or just a small number. Another issue for detecting proteins by mass spectrometry is high hydrophobicity (Lescuyer et al., 2004), for example, proteins that contain trans-membrane domains are often underrepresented in proteomic data sets. Finally, also factors like a low protein abundance, only context-specific expression, a high turnover rate or protein secretion might all hamper a successful detection of proteins (Elguoshy et al., 2016). Nonetheless, despite these barriers there are a few examples of translated overlapping genes with proteomic evidence. Notably, a large-scale study of 46 bacterial genomes found up to 261 cases of annotation ‘conflict’, i.e. overlaps greater than 40 base pairs with either proteomic evidence for both, or the unevidenced gene being annotated as something other than ‘hypothetical’ (Venter et al., 2011). A more recent study of 11 bacterial transcriptomes (Miravet-Verde et al., 2019) found 185 antisense transcripts previously annotated as non-coding could in fact code for proteins based on a random forest classifier (RanSEPs). A study in *Pseudomonas putida* found proteomic evidence for 44 ORFs embedded in antisense to annotated ORFs (Yang et al., 2016). An improved proteogenomics pipeline reported in a recent pre-print manuscript found numerous gene candidates in *Salmonella enterica* serovar Typhimurium, including a 199 amino acid long protein antisense to the annotated gene CBW18741 (Willems et al., 2019). It is interesting given the previous comment concerning the rate of phage to bacterial gene transfer that a BLAST search shows that this is likely a bacteriophage protein. The same study also found 18 antisense ORFs in *Deinoccocus radiodurans* supported by at least two peptides. A search in *Helicobacter pylori* mass spectrometry data from a previously published study designed to find small proteins (Müller et al., 2013) found evidence for a protein encoded by an ORF antisense to a proline/betaine transporter gene (Friedman et al., 2017). A recent discussion paper presented proteomic evidence for many small proteins (sORFs) and overlapping genes (‘altORFs’), but did not specifically consider antisense overlaps (Wu Orr et al. 2019).

Aside from proteomics datasets there is extensive publicly available high throughput RNA sequencing data which can be mined for further indicators of specific reproducible regulation of antisense ORFs. There are approximately 1500 relevant RNAseq studies from prokaryotes in the NCBI GEO database (Edgar et al., 2002), each with multiple samples; over 100 ribosome profiling studies, and a number of more bespoke methods which may also provide relevant information. CAPseq data, which discovers transcriptional start sites (Ettwiller et al., 2016), helps to delineate the borders of operons and their expression under different conditions. The new method SEnd-seq, through circularisation of transcripts, is able to detect both transcriptional start and termination sites with single nucleotide resolution (Ju et al., 2019). CHiPseq datasets indicate whether known transcription factors are associated with a particular operon of interest (Wade, 2015); other TF-binding assays also have potential for testing hypotheses concerning TF binding, e.g. DNAse footprinting (Haycocks and Grainger, 2016). Each of these methods is yet to be fully utilised in searching for the transcriptional regulation of overlapping genes. At the level of translation, there are a number of variations on ribosome profiling now available, including accurate prediction of translation initiation sites. The first study of ribosome profiling in bacteria used chloramphenicol in one of the two methods presented (Oh et al., 2011), which has since been shown to stall the ribosome at initiation and thereby can assist in inferring translation initiation site positions (Mohammad et al., 2019, Glaub et al., 2019). More precise stalling has been achieved with the use of tetracycline (Nakahigashi et al., 2016), retapamulin (Meydan et al., 2019), and the antibacterial peptide Onc112 (Weaver et al., 2019). Properties of ribosomes at different stages of translation, including initiation, have recently been studied in *Escherichia coli* K12 with TCP-seq (Sharma and Anand, 2019). Translation stop sites have also been specifically explored (Baggett et al., 2017). Most of these methods, outside the analysis of ribosome profiling discussed above, have not yet been applied to the detection or investigation of protein coding alternate frame ORFs, and any RNAs at these sites are assumed to be non-coding. Perhaps particularly useful will be ribosome profiling experiments conducted for cells grown in different conditions – many relevant contexts may however not be able to be surveyed due to technical limitations.

### Phenotypes of antisense proteins

An important indicator of functionality is specific regulation in response to defined environmental conditions. Some key canonical work in molecular genetics (e.g. (Jacob and Monod, 1961, Ames and Martin, 1964) has been concerned with the differential induction of genetic elements under varying environmental conditions. Specific differential induction is widely assumed in this kind of literature to be equivalent to function - how precisely to draw a line between functional and non-functional, given the inherent noisiness of biology, is not however entirely clear.

In general, what kind of phenotype is a good indicator of functionality? The most obvious case perhaps is an improvement in growth associated with expression of a genetic element. This could be either through improved growth following overexpression, or decreased growth following a deletion in the genomic sequence. Within an evolutionary context, a growth advantage effectively just is what it is to be ‘useful’ or ‘functional’. An example of this for antisense proteins is *citC*, discussed below. However, less intuitively, a decrease in growth associated with expression, as seen in the cases of *asa, laoB*, and *ano* is also an indicator of functionality in the right context. Most simply, the gene might literally function as a toxin. More generally though, overexpression of many functional genes is deleterious - in fact in *E. coli* the majority of annotated genes have a deleterious effect on growth in overexpression constructs (Kitagawa et al., 2005). Possible reasons for this high tendency towards deleterious phenotypes in bacteria compared with organisms such as yeast are discussed in (Bhattacharyya et al., 2016). This situation should perhaps not be surprising given the extreme optimality of bacterial metabolism (Schuetz et al., 2012); significant disturbance of such a finely tuned system is unlikely to be beneficial under most conditions. This general principle follows from, for instance, Fisher’s Geometric Model, in which random changes are less likely to be beneficial when a population is close to a fitness optimum (Tenaillon, 2014). Overexpression of a non-functional ‘junk’ genetic sequence however is also likely to be deleterious (Weisman and Eddy, 2017, Knopp and Andersson, 2018), so such a phenotype does not by itself provide evidence for functionality. What is important in the examples discussed above is that the deleterious growth phenotypes are observed as a significant difference between environmental (media) conditions. This implies a specificity of regulation which appears improbable under the ‘junk’ hypothesis, and so constitutes evidence of function.

A number of antisense overlapping genes in *E. coli* have been analysed regarding expression and phenotypes across different environmental conditions. The gene *nog1* is almost fully embedded in antisense to *citC*. A strand-specific deletion mutant has a growth advantage over the wildtype in LB, and a stronger advantage in medium supplemented with magnesium chloride (Fellner et al., 2015). The gene *asa*, embedded in antisense to a transcriptional regulator in *E. coli* O157:H7 strains, was found to be regulated in response to arginine, sodium, and different growth phases. Overexpression resulted in a negative growth phenotype in both excess sodium chloride and excess arginine, and no phenotype in LB medium (Vanderhaeghen et al., 2018). The gene *laoB* is embedded in antisense to a CadC-like transcriptional regulator (Figure 2A). A strand-specific genomic knock-out mutant was shown to provide a growth advantage specific to media supplemented with arginine. Further, the differential phenotype was replicated with the addition of inducible plasmid constructs bearing Δ*laoB* and WT *laoB*, showing that the phenotype is removed through complementation (Hücker et al., 2018b). How to mechanistically interpret such a growth advantage following gene knockout is unclear, but the condition-specific clear phenotype implies a functional role. The gene *ano* is nearly fully embedded antisense to an L,D-transpeptidase (Figure 2B). Similarly to *laoB*, a knock-out mutant showed a condition-specific phenotype. In this case it occurs in anaerobic conditions, and could be partially complemented with a plasmid construct (Hücker et al., 2018a). The putative protein-coding gene *aatS* was found in the pathogenic *E. coli* strain ETEC H10407 fully embedded in antisense to the ATP transporter ATB binding protein AatC (Haycocks and Grainger, 2016). It was shown to be transcribed, to have a functional ribosome binding sequence, and to have widespread homologs including a conserved domain of unknown function. Figure 2 illustrates the expression of three examples of antisense genes, with the gene in *Staphylococcus aureus* (Figure 2C) a special case, as discussed below.

**Figure 2:**
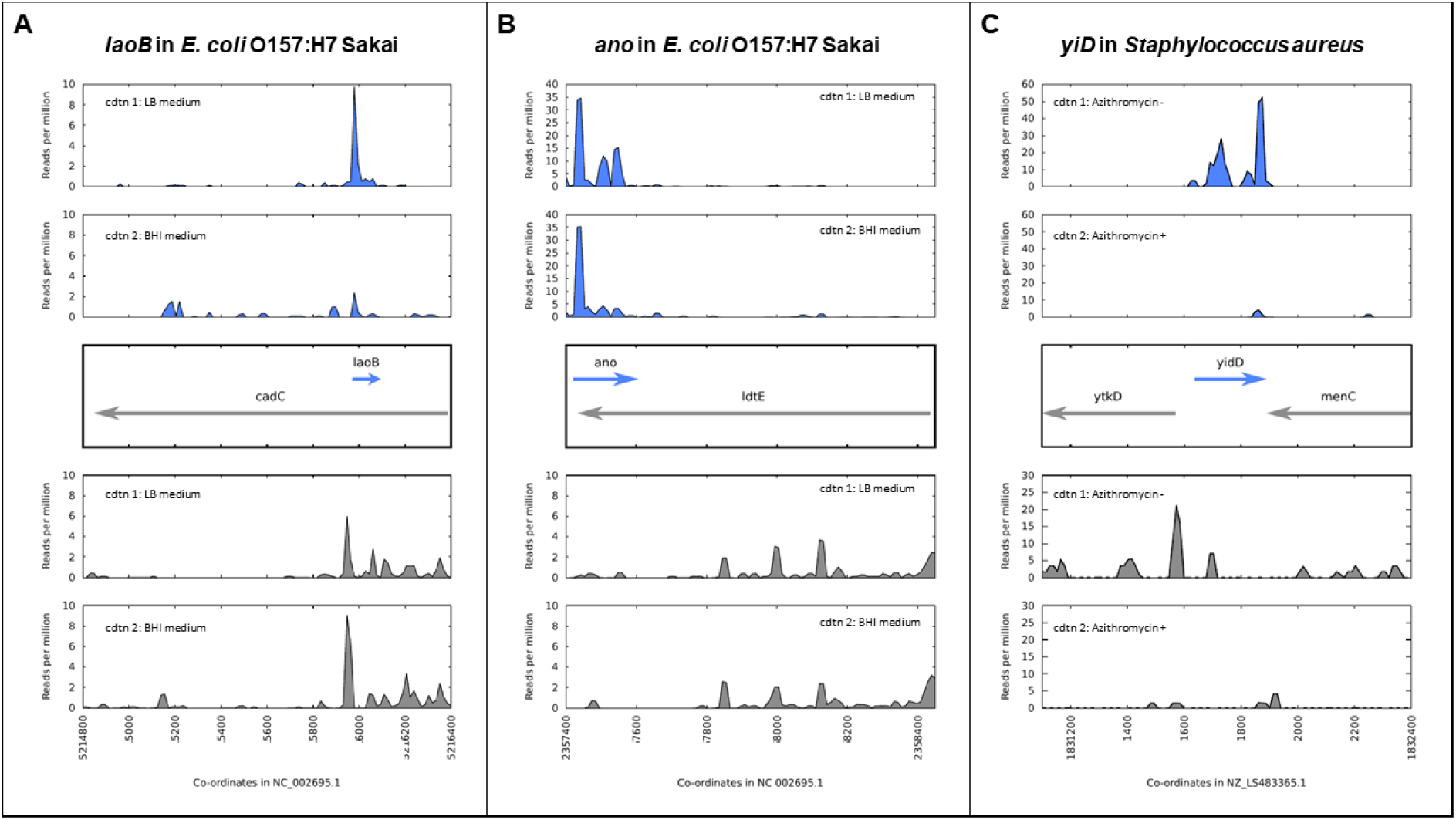
Expression and regulation of antisense genes as demonstrated by ribosome profiling experiments; aligned ribosome protected fragments shown as reads per million at each site after removal of rRNA and tRNA reads. A) *laoB* gene in O157:H7 Sakai; expression in LB medium is higher than in BHI. B) *ano* gene in E. coli O157:H7 Sakai; expression in LB medium versus BHI is constant. C) *yidD* (MW1733) gene, part of non-contiguous (antisense) operon, in *Staphylococcus aureus*. Expression in positive and negative strands is positively correlated - high in the absence of the antibiotic azithromycin, low when it is added.

Other than the high-level phenotypes (e.g. expression under particular conditions) determined for some candidates, very little is known about the possible roles or mechanisms of action of antisense proteins. Initial evidence (Vanderhaeghen, 2019) suggests that their cellular localisation may often be either secreted - similarly to the many unannotated intergenic protein-coding ORFs discovered in pathogenic strains of *E. coli*; (Hücker et al. 2017; Neuhaus et al. 2016). Signalling or interactions between cells will be a significant area to investigate regarding possible functions of antisense proteins. This suggestion is based on both the evidence gained so far for small proteins (Sberro et al., 2019) and the potential importance of any such genes. Any antisense proteins which are involved in signalling or inter-cell interactions may be of particular practical importance. This area is a crucial field of research as infectious disease continues to be a major health burden and the long natural history of interactions between microbes has been a fruitful source of new antimicrobial strategies.

### Simultaneous transcription?

In response to the evidence for overlapping genes, the question is often raised concerning how two genes could be simultaneously expressed from opposite strands. Indeed, the phenomenon of RNA polymerase collision is a real barrier to antisense transcription in at least some instances and is involved in transcriptional silencing or reduction via various mechanisms (Courtney and Chatterjee, 2014). Bypass of sense and antisense RNA polymerases has been shown for bacteriophage RNA polymerases (Ma and McAllister, 2009), but *in vitro* experiments have shown no such bypass in bacterial systems (Crampton et al., 2006). The role of accessory helicases in removing barriers to replication due to the presence of RNA polymerases has recently been highlighted (Hawkins et al., 2019), expanding on knowledge of simultaneous transcription and replication (Helmrich et al., 2013). It is conceivable that transcribing alongside the formation of a replication fork could facilitate antisense transcription, but this would restrict antisense transcription to the replication process. However, even in cases of collision of RNA polymerases operating in antisense, transcriptional stalling is not guaranteed. A recent study argues on the basis of simulations and careful assays with reporter constructs that RNA polymerases trailed by an active ribosome are, remarkably, about 13-times more likely to resume transcription following collision than those without the translation apparatus following (Hoffmann et al., 2019). This finding follows on from a range of similar work in recent years showing multiple mechanisms involved in ensuring that RNA polymerases stall and are subsequently released less in protein coding than non-coding RNAs (Proshkin et al., 2010, Brophy and Voigt, 2016, Ju et al., 2019). We suggest that this phenomenon likely applies to antisense embedded protein-coding genes as much as to convergent antisense transcripts and thereby facilitates antisense protein expression.

Recent detailed elucidation showed the working of an operon in *Staphylococcus aureus* with a functional gene encoded in antisense to a contiguous set of co-transcribed genes (Sáenz-Lahoya et al., 2019). The authors showed that despite being encoded on opposite strands (although not directly overlapping in this case), these elements comprised a single transcriptional unit (Figure 2C). This study highlights a mechanism which may be widespread and may apply to genes which are directly antiparallel as well. Results from Weaver et al. (2019) obtained by chromosomal tagging of three antisense proteins show that proteins encoded in antisense can be expressed simultaneously, i.e. under the same growth conditions. All this evidence for simultaneous antiparallel gene expression notwithstanding, it may be that antiparallel overlapping genes are generally translated under different conditions, or separated in time - this is yet to be determined.

### Evolution and constraint in antisense proteins

The evolutionary analysis of function at the nucleotide sequence level is a fairly recent development (Robinson-Rechavi, 2019), so we should not be surprised at unexpected results in this rapidly developing field. While the evolutionary analysis of antisense proteins in prokaryotes awaits further investigation of strong overlapping gene candidates, those discovered so far are typically relatively young (e.g. (Fellner et al., 2014, Fellner et al., 2015, Hücker et al., 2018a). This may be seen as a point against their functionality, particularly for candidates limited to just one species. However, a number of genome elements with undisputed functionality are also evolutionarily young. Various functional putatively ncRNA elements are known to have high evolutionary turnover, see e.g. Dutcher & Raghavan (2018). For instance, an sRNA found only in *E. coli* was shown to be derived from a pseudogenized bacteriophage gene (Kacharia et al., 2017). Also relevant here is the large literature on the functions of ‘orphan’ or taxonomically restricted genes restricted to a single genome or small clade (Satoshi and Nishikawa, 2004, Tautz and Domazet-Lošo, 2011) – and orphan genes may play diverse important roles in bacteria (Hu et al., 2009).

It appears likely that antisense proteins are often less constrained in sequence than most protein-coding genes currently known. For one, antisense proteins are typically quite small and hence unlikely to fold into complex structures. Secondly, given evidence that protein domains in overlapping genes may be situated so as to not overlap (Fernandes et al., 2016), it seems that overlapping gene sequences are unlikely to be comprised of a high proportion of constrained sequence domains. Many embedded open reading frames are actually well conserved beyond the genus though, for instance in the Enterobacteriaceae family. As a conservative example, when taking a subset of this family, the smallest clade including both *Citrobacter rodentium* and *Escherichia coli* (Figure 3A), we find that out of the 3391 antisense embedded ORFs predicted as having single homologs in all 13 representative genomes assessed, 29.5% exceed the conservation of the lower quartile of annotated genes (Figure 3B, 3C), as judged by median pairwise amino acid similarity between genomes.

**Figure 3:**
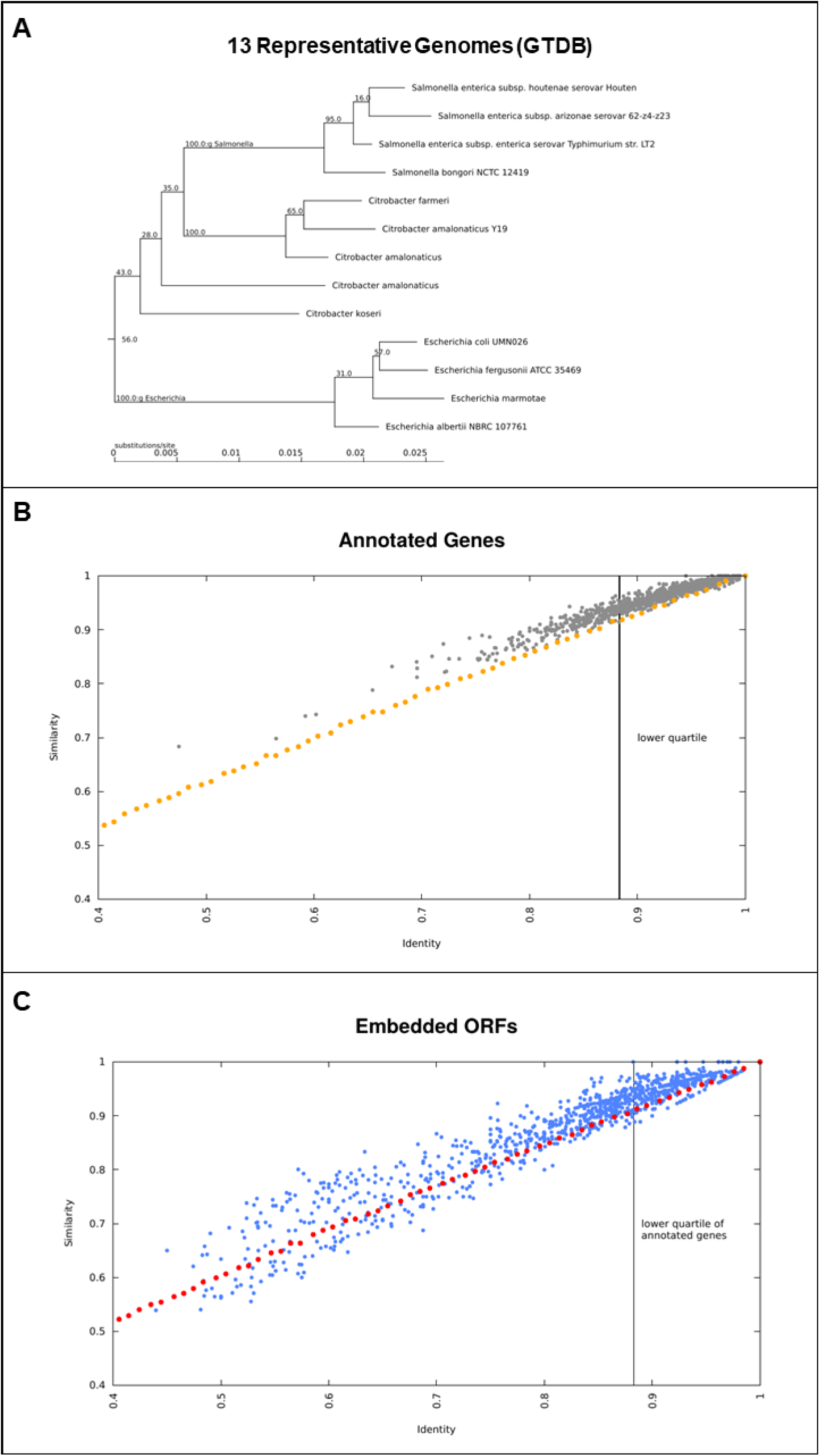
A subset of embedded antisense ORFs are well conserved in the genomes situated taxonomically between *E. coli* and *Citrobacter rodentium*. A) Phylogenetic tree showing 13 representative genomes used, derived from the genome taxonomy database (GTDB). B) median pairwise amino acid identity and similarity (grey) among orthologs for 1000 annotated genes with single copy orthologs in all of the 13 genomes. As a comparison, the effect on identity and similarity of adding random mutations to simulated sequences is shown (orange). There is a clear bias towards variants which result in higher ‘similarity’. C) conservation of antisense embedded ORFs (blue), as compared to the identity-similarity relationship observed for control randomly mutated sequences (red) simulated as before but translated in antisense. Many antisense embedded ORFs are highly conserved and a subset also shows a bias towards similarity.

Given the conservative nature of this analysis and that less than half of even annotated genes met the criterion of having single homologs in all these genomes, we posit that thousands of embedded antisense ORFs are sufficiently conserved beyond the *Escherichia* genus to be candidates for functional genes in this particular respect. Factors affecting these conservation statistics, and additional criteria for gene-likeness which distinguish coding from non-coding antisense sequences deserve further study.

The orange and red lines in Figure 3B and 3C show the effect of randomly mutating a sequence created based on the codon usage in the annotated genes in *E. coli* K12. The points represent median identities and similarities in comparison to originally simulated sequences, following successive rounds of random mutation, approximately mimicking the mutational distances observed between the orthologs of annotated genes and embedded ORFs. We suggest that two main results should be taken from Figure 3. Firstly, the blue cluster in the top right of Fig 3C shows that many embedded antisense ORFs are highly conserved across a significant evolutionary distance - they are not all immediately degraded following mutations in the alternate frame as might be naively assumed. Secondly, the bias above the orange and red lines shows that nearly all annotated genes and many embedded antisense ORFs tend towards fixing more similar mutations than might be predicted based on amino acid identity statistics alone. This result may be partly due to the structure of the genetic code, i.e. when ‘mother gene’ is conserved there is some tendency for conservation in the alternative strand (Wichmann and Ardern, 2019), but it is suggestive of a kind of purifying selection where mutations to biochemically similar amino acids are preferred in a subset of embedded antisense ORFs. It has previously been shown that long antisense ORFs appear more often in natural genomes than expected based on codon composition of annotated coding genes (Mir et al., 2012), another hint of selective processes preserving some antisense ORFs.

A recent, currently unpublished, study in *Mycobacterium tuberculosis* (Smith et al., 2019) has claimed that novel ORFs identified by ribosome profiling typically do not illustrate the strong codon bias evident in annotated mycobacterial genes and therefore cannot be expected to be functional. Given that many of these ORFs are situated in antisense to annotated genes, where the genetic code limits the possibilities for achieving optimal codon usage, this result is hardly surprising, and we suggest provides little evidence for the claim that they are nonfunctional. There is a problematic circularity here as well, as annotation of prokaryotic ORFs is based on models which take into account codon usage, based on usage in long ORFs - so short ORFs with ‘abnormal’ codon usage will likely remain unannotated, increasing reinforcing any bias in codon usage statistics in annotated genes. In general short and weakly expressed genes should not be expected to match ‘canonical’ highly expressed genes in terms of codon usage (Gupta and Ghosh, 2001), although the relationship between expression and codon usage is not straightforward (Dos Reis et al., 2003). Careful evolutionary sequence analyses are required here. A study of some putative same-strand overlapping genes also suggested that they are not under constraint (Meydan et al., 2019). However, more biologically nuanced analyses of sequence constraint, for instance after partitioning the homologs into phylostrata, would be useful. Further, a fundamental assumption of methods for detecting selection, e.g. Firth (2014), Wei & Zhang (2015), is the neutrality of synonymous mutations, but this assumption has been shown to be false, with the rate of synonymous mutations varying widely across sites (Wisotsky et al., 2020). The extent to which this affects conclusions regarding dN/dS as calculated with the various available methods remains unexplored. More generally, to our knowledge, there has been no demonstration of any synthesis of non-functional protein in prokaryotes. The high bioenergetic cost of protein production (Lynch and Marinov, 2015) would seem to militate against such a phenomenon being widespread in bacteria, where costs are minimised through gene loss (Koskiniemi et al., 2012). As such we argue that the default assumption following demonstration of a clear signal of translation should be that the product plays a functional role.

### The context: unexpected complexity

The historical trajectory in bacterial genomics has been towards finding previously unappreciated layers of complexity (Grainger, 2016). In particular, the number of different kinds of functional elements recognised has continued to increase in recent years. Examples of genetic elements previously ignored or written off as background noise which are now known to be functional in some or many cases include antisense transcription, small RNAs and microRNAs, proteins with alternative start sites, small proteins (Storz et al., 2014), and micropeptides. Antisense transcription has been widely disregarded as noise (Lloréns-Rico et al., 2016, Raghavan et al., 2012). However, despite these generalisations, these elements have recently been found to at least sometimes have physiological roles (Wade and Grainger, 2014, Lejars et al., 2019). Functional RNAs which are not yet well understood include structured noncoding RNAs such as riboswitches (Stav et al., 2019, Hücker et al., 2017). Proteins with alternative start sites, designated isoforms or ‘proteoforms’, have also been reported in a few bacterial systems (Berry et al., 2016, Nakahigashi et al., 2016, Meydan et al., 2019). These genetic elements are all yet to be incorporated into genome annotation files and gene prediction algorithms. As such, genome annotation is years behind the leading edge of research in bacterial genetics, and various functional elements remain unannotated.

### Recommendations for further research

Even recent attempts at comprehensive studies of small proteins have tended to ignore antisense proteins or to use methods unintentionally biased against them - perhaps unsurprising given the reigning paradigm in genome annotation, which excludes substantive overlaps as a matter of principle. As an example, the NCBI prokaryotic genome annotation standards include among the minimum standards that there can be “[n]o gene completely contained in another gene on the same or opposite strand” (NCBI, 2020). For instance, a recent study investigated small proteins in the human microbiome (Sberro et al., 2019), finding hundreds of previously unknown small proteins with evidence from evolutionary sequence constraint, and many also with evidence of transcription and/or translation. Two key steps were the use of MetaProdigal for gene prediction and RNAcode for inference of conservation. Both of these are implicitly biased against overlapping genes, in that Prodigal explicitly excludes long overlaps, and RNAcode looks for patterns of sequence constraint associated with normal non-overlapping genes, which are unlikely to be found in overlapping genes.

The bacteriological research community ought to relinquish the common assumption that unannotated functional elements are only to be found in intergenic regions. We must also be aware that antisense regions often need to be treated differently from intergenic regions, for instance in analyses of sequence constraint. Developing appropriate corrections to take into account the sequence context of antisense overlapping ORFs is an important area for further work. A major emphasis should be on high-throughput functional studies. For in-depth laboratory studies dissecting the details of an overlapping gene’s regulation and function, the focus should be on the strongest candidates as determined with sequence and expression data. One key criterion here is evidence of reproducible regulated translation from one of the various ribosome profiling methods now available. Sequence properties determined from such sets should help to find strong candidates which are not expressed under already-assayed conditions. It is also clear that further advances in proteomics for small proteins should result in proteomic evidence for the translation of many more antisense proteins in bacteria and other systems. Following on from this, the discovery of any protein structures would be a major step forward towards understanding the molecular mechanisms of function. Finally, studying the evolutionary history of antisense proteins may provide useful insights on function. In this aspect these genes have a significant advantage over others in that their genomic context is relatively fixed by the gene in which they are embedded. This study has focused on eubacteria, but the same principles conceivably apply in archaea. A recent study, for instance, chose to only consider same-frame overlapping ORFs (proteoforms) on account of an absence of proteomics results and reliable BLAST hits for out-of-frame overlapping ORFs (Ten-Caten et al., 2018). Neither of these negative results are surprising however given the limitations of proteomics discussed above and the current bias against annotating out-of-frame overlaps; as such, archaeal datasets ought also be re-examined for functional overlapping genes.

In summary, what is required in order to assign the descriptor ‘functional/ to a putative gene, such as a gene encoded in antisense to a known gene? Regarding evolutionary evidence, a codon-level pattern of sequence constraint is sufficient to guarantee function, as constraint matching expectations for amino acids is unexpected in coding sequences. Detecting such constraint is a challenge for antisense sequences however. Regarding evidence from wet lab experiments, a condition-specific phenotype is also sufficient to establish functionality. The ‘gold standard’ in this area would be a condition-specific negative growth phenotype in a genomic knock-out mutant, which could be complemented in *trans* (e.g. with a plasmid construct). Regarding high-throughput evidence, significant protein expression is evidence of functionality in highly optimized bacterial genomes, particularly if shown to be consistent across species or highly diverged strains. Appropriate thresholds for significant expression and sufficient evolutionary divergence in order to be able to confidently infer function are yet to be established. While each of these three lines of evidence is arguably sufficient to establish function, none is necessary, as there are functional elements which fail to meet at least one of these criteria.

We have collated evidence from diverse bacteria (including the genera *Escherichia, Pseudomonas*, and *Mycobacterium*) for protein coding ORFs embedded in antisense to annotated genes, discussed reasons to believe that they are biologically functional, and responded to common objections, informed by the most recent work in bacterial molecular genetics. We suggest that a pro-function attitude regarding antisense prokaryotic transcripts and the antisense translatome is both more useful for research and justified by multiple lines of evidence. How many of these elements are functional and what they do remain contentious however, and worthy of significant further investigation.

## Methods

For Figure 1, positions of previously discovered putative antiparallel genes in *Escherichia coli* K12 and *Mycobacterium tuberculosis* were extracted from the supplementary data of previous studies. (Weaver et al., 2019, Friedman et al., 2017, Smith et al., 2019); information on those with ribosome profiling reads was provided by Robin Friedman. Positions are shown visualised with Circos (Krzywinski et al., 2009).

For Figure 2, ribosome profiling (“RIBO-seq”) data was visualised to show examples of antisense overlapping genes. In each case, adapter sequences were predicted using DNApi.py (Tsuji and Weng, 2016), trimmed with cutadapt (Martin, 2011) using a minimum length of 19 and quality score of 10, and aligned with bowtie2 (Langmead and Salzberg, 2012). Fastq data from SRR5874479 (LB) and SRR5874484 (BHI) for *E. coli* O157:H7 Sakai was aligned against the genome GCF_000008865.1_ASM886v1. Fastq data from SRR1265839 (azi minus) and SRR1265836 (azi plus) for *Staphylococcus aureus* was aligned against genome GCF_900475245.1_43024_E01. Reads mapping at each site per million total mapped reads (RPM) are calculated from aligned bam files with reads mapping to rRNA and tRNA locations removed, using samtools (Li et al., 2009). Images of RPM in the region around the putative antisense gene are drawn in gnuplot with ‘smooth csplines’.

For Figure 3 the relationship between similarity and identity in comparisons of different ORF homologs was compared. Representative genomes from release 89 of the genome taxonomy database (GTDB) (Parks et al., 2018) in the smallest clade uniting *E. coli* and *Citrobacter rodentium* (Fig 3A) were chosen. Of these 23 strains, 13 had a GenBank genome and feature table with the same accession version available. These were downloaded, and annotated ORFs in each compared to each other using OrthoFinder (Emms and Kelly, 2015). Genes with a single copy ortholog present in all 13 genomes were extracted, and members of each ortholog family were aligned against each other using the EMBOSS (Rice et al., 2000) program needleall to determine median similarity and identity at the amino acid level (Figure 3B). As a control, 50 sequences of 333 codons length were created based on codon usage in *E. coli* K12, using EMBOSS programs cusp and makenucseq. These were then mutated through 70 rounds of point mutation (10 mutations per round) using the EMBOSS program msbar, and translated in order to determine the relationship between varying levels of amino acid identity and similarity. In each case the mutated sequences were compared to the original simulated sequence they were derived from, using EMBOSS needle. For each percent decrease in identity observed, results were collated and the median values of identity and similarity reported. The procedure used initially for annotated genes was repeated using all antisense embedded ORFs, found using a Perl script and Bedtools (Quinlan and Hall, 2010). A negative control for these sequences was also created similarly to before, but using an antisense reading frame. As the particular antisense frame used had no significant effect on the sequence similarities obtained in the simulation, for the data shown the initial sequences based on codon usage in *E. coli* K12 were directly reverse complemented with no further frame-shift, prior to the 70 rounds of random mutation.

## Acknowledgements

Thanks to Christina Ludwig for advice on proteomics references, Christopher Huptas for development of an ORF finder Perl script, and Robin Friedman for providing ribosomal profiling information for putative sRNAs.

## References

Affram, Y., Zapata, J. C., Zhou, W., Pazgier, M., Iglesias-Ussel, M., Ray, K., Latinovic, O. & Romerio, F. 2019. PJ-1 The HIV-1 antisense protein ASP is a structural protein of the viral envelope. JAIDS Journal of Acquired Immune Deficiency Syndromes, 81, 79.

Ames, B. N. & Martin, R. G. 1964. Biochemical aspects of genetics: the operon. Annual review of biochemistry, 33, 235–258.

Ardern, Z. 2018. Dysfunction, disease, and the limits of selection. Biological Theory, 13, 4–9.

Baggett, N. E., Zhang, Y. & Gross, C. A. 2017. Global analysis of translation termination in E. coli. PLoS genetics, 13, e1006676.

Barrell, B. G., Air, G. & Hutchison, C. 1976. Overlapping genes in bacteriophage φX174. Nature, 264, 34–41.

Bendall, M. L., Stevens, S. L., Chan, L.-K., Malfatti, S., Schwientek, P., Tremblay, J., Schackwitz, W., Martin, J., Pati, A. & Bushnell, B. 2016. Genome-wide selective sweeps and gene-specific sweeps in natural bacterial populations. The ISME journal, 10, 1589–1601.

Berry, I. J., Steele, J. R., Padula, M. P. & Djordjevic, S. P. 2016. The application of terminomics for the identification of protein start sites and proteoforms in bacteria. Proteomics, 16, 257–272.

Bhattacharyya, S., Bershtein, S., Yan, J., Argun, T., Gilson, A. I., Trauger, S. A. & Shakhnovich, E. I. 2016. Transient protein-protein interactions perturb E. coli metabolome and cause gene dosage toxicity. Elife, 5, e20309.

Brandon, R. N. 2013. A general case for functional pluralism. Functions: Selection and mechanisms. Springer.

Brophy, J. A. & Voigt, C. A. 2016. Antisense transcription as a tool to tune gene expression. Molecular systems biology, 12.

Cassan, E., Arigon-Chifolleau, A. M., Mesnard, J. M., Gross, A. & Gascuel, O. 2016. Concomitant emergence of the antisense protein gene of HIV-1 and of the pandemic. Proc Natl Acad Sci U S A, 113, 11537–11542.

Cheetham, S. W., Faulkner, G. J. & Dinger, M. E. 2019. Overcoming challenges and dogmas to understand the functions of pseudogenes. Nature Reviews Genetics, 1–11.

Chou, K.-C., Zhang, C.-T. & Elrod, D. W. 1996. Do “antisense proteins” exist? Journal of protein chemistry, 15, 59–61.

Courtney, C. & Chatterjee, A. 2014. cis-Antisense RNA and transcriptional interference: coupled layers of gene regulation. J. Gene Ther, 2, 1–9.

Crampton, N., Bonass, W. A., Kirkham, J., Rivetti, C. & Thomson, N. H. 2006. Collision events between RNA polymerases in convergent transcription studied by atomic force microscopy. Nucleic acids research, 34, 5416–5425.

Doolittle, W. F. 2018. We simply cannot go on being so vague about ‘function’. Genome biology, 19, 1–3.

Doolittle, W. F., Brunet, T. D., Linquist, S. & Gregory, T. R. 2014. Distinguishing between “function” and “effect” in genome biology. Genome biology and evolution, 6, 1234–1237.

Dos Reis, M., Wernisch, L. & Savva, R. 2003. Unexpected correlations between gene expression and codon usage bias from microarray data for the whole Escherichia coli K-12 genome. Nucleic acids research, 31, 6976–6985.

Dutcher, H. A. & Raghavan, R. 2018. Origin, evolution, and loss of bacterial small RNAs. Regulating with RNA in Bacteria and Archaea, 487–497.

Edgar, R., Domrachev, M. & Lash, A. E. 2002. Gene Expression Omnibus: NCBI gene expression and hybridization array data repository. Nucleic acids research, 30, 207–210.

Elguoshy, A., Magdeldin, S., Xu, B., Hirao, Y., Zhang, Y., Kinoshita, N., Takisawa, Y., Nameta, M., Yamamoto, K. & El-Refy, A. 2016. Why are they missing: Bioinformatics characterization of missing human proteins. Journal of proteomics, 149, 7–14.

Ellis, J. C. & Brown, J. W. 2003. Genes within genes within bacteria. Trends in biochemical sciences, 28, 521–523.

Emms, D. M. & Kelly, S. 2015. OrthoFinder: solving fundamental biases in whole genome comparisons dramatically improves orthogroup inference accuracy. Genome biology, 16, 157.

Ettwiller, L., Buswell, J., Yigit, E. & Schildkraut, I. 2016. A novel enrichment strategy reveals unprecedented number of novel transcription start sites at single base resolution in a model prokaryote and the gut microbiome. BMC Genomics, 17, 199.

Fellner, L., Bechtel, N., Witting, M. A., Simon, S., Schmitt-Kopplin, P., Keim, D., Scherer, S. & Neuhaus, K. 2014. Phenotype of htgA (mbiA), a recently evolved orphan gene of Escherichia coli and Shigella, completely overlapping in antisense to yaaW. FEMS Microbiol Lett, 350, 57–64.

Fellner, L., Simon, S., Scherling, C., Witting, M., Schober, S., Polte, C., Schmitt-Kopplin, P., Keim, D. A., Scherer, S. & Neuhaus, K. 2015. Evidence for the recent origin of a bacterial protein-coding, overlapping orphan gene by evolutionary overprinting. BMC Evol Biol, 15, 283.

Fernandes, J. D., Faust, T. B., Strauli, N. B., Smith, C., Crosby, D. C., Nakamura, R. L., Hernandez, R. D. & Frankel, A. D. 2016. Functional segregation of overlapping genes in HIV. Cell, 167, 1762-1773. e12.

Firth, A. E. 2014. Mapping overlapping functional elements embedded within the protein-coding regions of RNA viruses. Nucleic acids research, 42, 12425–12439.

Friedman, R. C., Kalkhof, S., Doppelt-Azeroual, O., Mueller, S. A., Chovancova, M., Von Bergen, M. & Schwikowski, B. 2017. Common and phylogenetically widespread coding for peptides by bacterial small RNAs. BMC genomics, 18, 553.

Georg, J. & Hess, W. R. 2018. Widespread antisense transcription in prokaryotes. Regulating with RNA in Bacteria and Archaea, 191–210.

Gibson, B. & Eyre-Walker, A. 2019. Investigating evolutionary rate variation in bacteria. Journal of molecular evolution, 87, 317–326.

Gimpel, M. & Brantl, S. 2017. Dual-function small regulatory RNAs in bacteria. Molecular microbiology, 103, 387–397.

Glaub, A. S., Huptas, C., Neuhaus, K. & Ardern, Z. 2019. Improving Bacterial Ribosome Profiling Data Quality. bioRxiv, 863266.

Goodhead, I. & Darby, A. C. 2015. Taking the pseudo out of pseudogenes. Current opinion in microbiology, 23, 102–109.

Grainger, D. C. 2016. The unexpected complexity of bacterial genomes. Microbiology, 162, 1167–1172.

Graur, D., Zheng, Y. & Azevedo, R. B. 2015. An evolutionary classification of genomic function. Genome biology and evolution, 7, 642–645.

Graur, D., Zheng, Y., Price, N., Azevedo, R. B., Zufall, R. A. & Elhaik, E. 2013. On the immortality of television sets:”function” in the human genome according to the evolution-free gospel of ENCODE. Genome biology and evolution, 5, 578–590.

Gupta, S. & Ghosh, T. 2001. Gene expressivity is the main factor in dictating the codon usage variation among the genes in Pseudomonas aeruginosa. Gene, 273, 63–70.

Harrison, E. & Brockhurst, M. A. 2017. Ecological and evolutionary benefits of temperate phage: what does or doesn’t kill you makes you stronger. BioEssays, 39, 1700112.

Hawkins, M., Dimude, J. U., Howard, J. A. L., Smith, A. J., Dillingham, M. S., Savery, N. J., Rudolph, C. J. & Mcglynn, P. 2019. Direct removal of RNA polymerase barriers to replication by accessory replicative helicases. Nucleic acids research, 47, 5100–5113.

Haycocks, J. R. & Grainger, D. C. 2016. Unusually situated binding sites for bacterial transcription factors can have hidden functionality. PloS one, 11.

Helmrich, A., Ballarino, M., Nudler, E. & Tora, L. 2013. Transcription-replication encounters, consequences and genomic instability. Nature structural & molecular biology, 20, 412–418.

Hoffmann, S. A., Hao, N., Shearwin, K. E. & Arndt, K. M. 2019. Characterizing transcriptional interference between converging genes in bacteria. ACS synthetic biology, 8, 466–473.

Hu, P., Janga, S. C., Babu, M., Dίaz-Mejίa, J. J., Butland, G., Yang, W., Pogoutse, O., Guo, X., Phanse, S. & Wong, P. 2009. Global functional atlas of Escherichia coli encompassing previously uncharacterized proteins. PLoS biology, 7.

Hücker, S. M., Simon, S., Scherer, S. & Neuhaus, K. 2017. Transcriptional and translational regulation by RNA thermometers, riboswitches and the sRNA DsrA in Escherichia coli O157: H7 Sakai under combined cold and osmotic stress adaptation. FEMS microbiology letters, 364.

Hücker, S. M., Vanderhaeghen, S., Abellan-Schneyder, I., Scherer, S. & Neuhaus, K. 2018a. The Novel Anaerobiosis-Responsive Overlapping Gene ano Is Overlapping Antisense to the Annotated Gene ECs2385 of Escherichia coli O157:H7 Sakai. Frontiers in Microbiology, 9.

Hücker, S. M., Vanderhaeghen, S., Abellan-Schneyder, I., Wecko, R., Simon, S., Scherer, S. & Neuhaus, K. 2018b. A novel short L-arginine responsive protein-coding gene (laoB) antiparallel overlapping to a CadC-like transcriptional regulator in Escherichia coli O157:H7 Sakai originated by overprinting. BMC Evol Biol, 18, 21.

Ingolia, N. T., Ghaemmaghami, S., Newman, J. R. & Weissman, J. S. 2009. Genome-wide analysis in vivo of translation with nucleotide resolution using ribosome profiling. science, 324, 218–223.

Jacob, F. & Monod, J. 1961. Genetic regulatory mechanisms in the synthesis of proteins. Journal of molecular biology, 3, 318–356.

Jeong, Y., Kim, J.-N., Kim, M. W., Bucca, G., Cho, S., Yoon, Y. J., Kim, B.-G., Roe, J.-H., Kim, S. C. & Smith, C. P. 2016. The dynamic transcriptional and translational landscape of the model antibiotic producer Streptomyces coelicolor A3 (2). Nature communications, 7, 11605.

Ju, X., Li, D. & Liu, S. 2019. Full-length RNA profiling reveals pervasive bidirectional transcription terminators in bacteria. Nature microbiology, 4, 1907–1918.

Kacharia, F. R., Millar, J. A. & Raghavan, R. 2017. Emergence of new sRNAs in enteric bacteria is associated with low expression and rapid evolution. Journal of molecular evolution, 84, 204–213.

Keeling, D. M., Garza, P., Nartey, C. M. & Carvunis, A.-R. 2019. The meanings of’function’in biology and the problematic case of de novo gene emergence. eLife, 8.

Kitagawa, M., Ara, T., Arifuzzaman, M., Ioka-Nakamichi, T., Inamoto, E., Toyonaga, H. & Mori, H. 2005. Complete set of ORF clones of Escherichia coli ASKA library (A Complete S et of E. coli K-12 ORF A rchive): Unique Resources for Biological Research. DNA research, 12, 291–299.

Knopp, M. & Andersson, D. I. 2018. No beneficial fitness effects of random peptides. Nature ecology & evolution, 2, 1046–1047.

Koskiniemi, S., Sun, S., Berg, O. G. & Andersson, D. I. 2012. Selection-driven gene loss in bacteria. PLoS genetics, 8.

Krzywinski, M., Schein, J., Birol, I., Connors, J., Gascoyne, R., Horsman, D., Jones, S. J. & Marra, M. A. 2009. Circos: an information aesthetic for comparative genomics. Genome research, 19, 1639–1645.

Langmead, B. & Salzberg, S. L. 2012. Fast gapped-read alignment with Bowtie 2. Nature methods, 9, 357–359.

Lejars, M., Kobayashi, A. & Hajnsdorf, E. 2019. Physiological roles of antisense RNAs in prokaryotes. Biochimie.

Lescuyer, P., Hochstrasser, D. F. & Sanchez, J. C. 2004. Comprehensive proteome analysis by chromatographic protein prefractionation. Electrophoresis, 25, 1125–1135.

Li, H., Handsaker, B., Wysoker, A., Fennell, T., Ruan, J., Homer, N., Marth, G., Abecasis, G. & Durbin, R. 2009. The sequence alignment/map format and SAMtools. Bioinformatics, 25, 2078–2079.

Lloréns-Rico, V., Cano, J., Kamminga, T., Gil, R., Latorre, A., Chen, W.-H., Bork, P., Glass, J. I., Serrano, L. & Lluch-Senar, M. 2016. Bacterial antisense RNAs are mainly the product of transcriptional noise. Science advances, 2, e1501363.

Lynch, M. & Marinov, G. K. 2015. The bioenergetic costs of a gene. Proc Natl Acad Sci U S A, 112, 15690–5.

Ma, N. & Mcallister, W. T. 2009. In a head-on collision, two RNA polymerases approaching one another on the same DNA may pass by one another. Journal of molecular biology, 391, 808–812.

Martin, M. 2011. Cutadapt removes adapter sequences from high-throughput sequencing reads. EMBnet. journal, 17, 10–12.

Meydan, S., Marks, J., Klepacki, D., Sharma, V., Baranov, P. V., Firth, A. E., Margus, T., Kefi, A., Vazquez-Laslop, N. & Mankin, A. S. 2019. Retapamulin-assisted ribosome profiling reveals the alternative bacterial proteome. Molecular cell, 74, 481-493. e6.

Mir, K., Neuhaus, K., Scherer, S., Bossert, M. & Schober, S. 2012. Predicting statistical properties of open reading frames in bacterial genomes. PLoS One, 7.

Miravet-Verde, S., Ferrar, T., Espadas-GarcίA, G., Mazzolini, R., Gharrab, A., Sabido, E., Serrano, L. & Lluch-Senar, M. 2019. Unraveling the hidden universe of small proteins in bacterial genomes. Molecular systems biology, 15.

Mohammad, F., Green, R. & Buskirk, A. R. 2019. A systematically-revised ribosome profiling method for bacteria reveals pauses at single-codon resolution. Elife, 8, e42591.

Müller, S. A., Findeiß, S., Pernitzsch, S. R., Wissenbach, D. K., Stadler, P. F., Hofacker, I. L., Von Bergen, M. & Kalkhof, S. 2013. Identification of new protein coding sequences and signal peptidase cleavage sites of Helicobacter pylori strain 26695 by proteogenomics. Journal of proteomics, 86, 27–42.

Nakahigashi, K., Takai, Y., Kimura, M., Abe, N., Nakayashiki, T., Shiwa, Y., Yoshikawa, H., Wanner, B. L., Ishihama, Y. & Mori, H. 2016. Comprehensive identification of translation start sites by tetracycline-inhibited ribosome profiling. DNA Research, 23, 193–201.

NCBI. 2020. NCBI Prokaryotic Genome Annotation Standards [Online]. [Online]. National Center for Biotechnology Information, U.S. National Library of Medicine Available: https://www.ncbi.nlm.nih.gov/genome/annotation_prok/standards/ [Accessed 20.02.2020].

Nelson, C. W., Ardern, Z. & Wei, X. 2019. OLGenie: Estimating Natural Selection to Predict Functional Overlapping Genes. bioRxiv.

Neuhaus, K., Landstorfer, R., Simon, S., Schober, S., Wright, P. R., Smith, C., Backofen, R., Wecko, R., Keim, D. A. & Scherer, S. 2017. Differentiation of ncRNAs from small mRNAs in Escherichia coli O157: H7 EDL933 (EHEC) by combined RNAseq and RIBOseq–ryhB encodes the regulatory RNA RyhB and a peptide, RyhP. BMC genomics, 18, 216.

Oh, E., Becker, A. H., Sandikci, A., Huber, D., Chaba, R., Gloge, F., Nichols, R. J., Typas, A., Gross, C. A., Kramer, G., Weissman, J. S. & Bukau, B. 2011. Selective ribosome profiling reveals the cotranslational chaperone action of trigger factor in vivo. Cell, 147, 1295–308.

Owen, S. V., Canals, R., Wenner, N., Hammarlöf, D. L., Kröger, C. & Hinton, J. C. 2019. A window into lysogeny: Revealing temperate phage biology with transcriptomics. BioRxiv, 787010.

Parks, D. H., Chuvochina, M., Waite, D. W., Rinke, C., Skarshewski, A., Chaumeil, P.-A. & Hugenholtz, P. 2018. A standardized bacterial taxonomy based on genome phylogeny substantially revises the tree of life. Nature biotechnology, 36, 996–1004.

Pavesi, A. 2019. Asymmetric evolution in viral overlapping genes is a source of selective protein adaptation. Virology, 532, 39–47.

Pavesi, A., Vianelli, A., Chirico, N., Bao, Y., Blinkova, O., Belshaw, R., Firth, A. & Karlin, D. 2018. Overlapping genes and the proteins they encode differ significantly in their sequence composition from non-overlapping genes. PloS one, 13.

Proshkin, S., Rahmouni, A. R., Mironov, A. & Nudler, E. 2010. Cooperation between translating ribosomes and RNA polymerase in transcription elongation. Science, 328, 504–508.

Quinlan, A. R. & Hall, I. M. 2010. BEDTools: a flexible suite of utilities for comparing genomic features. Bioinformatics, 26, 841–842.

Raghavan, R., Sloan, D. B. & Ochman, H. 2012. Antisense transcription is pervasive but rarely conserved in enteric bacteria. MBio, 3, e00156–12.

Rice, P., Longden, I. & Bleasby, A. 2000. EMBOSS: the European molecular biology open software suite. Elsevier current trends.

Robinson-Rechavi, M. 2019. Molecular evolution and gene function. arXiv preprint arXiv:1910.01940.

Sáenz-Lahoya, S., Bitarte, N., Garcίa, B., Burgui, S., Vergara-Irigaray, M., Valle, J., Solano, C., Toledo-Arana, A. & Lasa, I. 2019. Noncontiguous operon is a genetic organization for coordinating bacterial gene expression. Proceedings of the National Academy of Sciences, 116, 1733–1738.

Saha, D., Podder, S., Panda, A. & Ghosh, T. C. 2016. Overlapping genes: a significant genomic correlate of prokaryotic growth rates. Gene, 582, 143–147.

Satoshi, F. & Nishikawa, K. 2004. Estimation of the number of authentic orphan genes in bacterial genomes. DNA research, 11, 219–231.

Sberro, H., Fremin, B. J., Zlitni, S., Edfors, F., Greenfield, N., Snyder, M. P., Pavlopoulos, G. A., Kyrpides, N. C. & Bhatt, A. S. 2019. Large-scale analyses of human microbiomes reveal thousands of small, novel genes. Cell, 178, 1245-1259. e14.

Schuetz, R., Zamboni, N., Zampieri, M., Heinemann, M. & Sauer, U. 2012. Multidimensional optimality of microbial metabolism. Science, 336, 601–604.

Sela, I., Wolf, Y. I. & Koonin, E. V. 2019. Selection and genome plasticity as the key factors in the evolution of bacteria. Physical Review X, 9, 031018.

Sharma, H. & Anand, B. 2019. Ribosome assembly defects subvert initiation Factor3 mediated scrutiny of bona fide start signal. Nucleic acids research, 47, 11368–11386.

Smith, C., Canestrari, J., Wang, J., Derbyshire, K., Gray, T. & Wade, J. 2019. Pervasive Translation in Mycobacterium tuberculosis. bioRxiv, 665208.

Stav, S., Atilho, R. M., Arachchilage, G. M., Nguyen, G., Higgs, G. & Breaker, R. R. 2019. Genome-wide discovery of structured noncoding RNAs in bacteria. BMC microbiology, 19, 66.

Storz, G., Wolf, Y. I. & Ramamurthi, K. S. 2014. Small proteins can no longer be ignored. Annu Rev Biochem, 83, 753–77.

Takeuchi, N., Cordero, O. X., Koonin, E. V. & Kaneko, K. 2015. Gene-specific selective sweeps in bacteria and archaea caused by negative frequency-dependent selection. BMC biology, 13, 20.

Tautz, D. & Domazet-Lošo, T. 2011. The evolutionary origin of orphan genes. Nature Reviews Genetics, 12, 692–702.

Ten-Caten, F., Vêncio, R. Z., Lorenzetti, A. P. R., Zaramela, L. S., Santana, A. C. & Koide, T. 2018. Internal RNAs overlapping coding sequences can drive the production of alternative proteins in archaea. RNA biology, 15, 1119–1132.

Tenaillon, O. 2014. The utility of Fisher’s geometric model in evolutionary genetics. Annual review of ecology, evolution, and systematics, 45, 179–201.

Tsuji, J. & Weng, Z. 2016. DNApi: a de novo adapter prediction algorithm for small RNA sequencing data. PloS one, 11.

Vanderhaeghen, S. 2019. Overlapping genes in E. coli EDL933 (EHEC) - Phylostratigraphy of alternative reading frames and functional analysis of the candidate gene asa. PhD, Technischen Universität München.

Vanderhaeghen, S., Zehentner, B., Scherer, S., Neuhaus, K. & Ardern, Z. 2018. The novel EHEC gene asa overlaps the TEGT transporter gene in antisense and is regulated by NaCl and growth phase. Scientific reports, 8, 17875.

Venter, E., Smith, R. D. & Payne, S. H. 2011. Proteogenomic analysis of bacteria and archaea: a 46 organism case study. PloS one, 6.

Wade, J. T. 2015. Mapping transcription regulatory networks with ChIP-seq and RNA-seq. Prokaryotic systems biology. Springer.

Wade, J. T. & Grainger, D. C. 2014. Pervasive transcription: illuminating the dark matter of bacterial transcriptomes. Nature Reviews Microbiology, 12, 647–653.

Wadler, C. S. & Vanderpool, C. K. 2007. A dual function for a bacterial small RNA: SgrS performs base pairing-dependent regulation and encodes a functional polypeptide. Proceedings of the National Academy of Sciences, 104, 20454–20459.

Weaver, J., Mohammad, F., Buskirk, A. R. & Storz, G. 2019. Identifying small proteins by ribosome profiling with stalled initiation complexes. MBio, 10, e02819–18.

Wei, X. & Zhang, J. 2015. A simple method for estimating the strength of natural selection on overlapping genes. Genome biology and evolution, 7, 381–390.

Weisman, C. M. & Eddy, S. R. 2017. Gene Evolution: Getting Something from Nothing. Curr Biol, 27, R661–R663.

Wichmann, S. & Ardern, Z. 2019. Optimality in the standard genetic code is robust with respect to comparison code sets. Biosystems, 185, 104023.

Willems, P., Fijalkowski, I. & Van Damme, P. 2019. Lost and found: re-searching and re-scoring proteomics data aids the discovery of bacterial proteins and improves proteome coverage. bioRxiv.

Willis, S. & Masel, J. 2018. Gene birth contributes to structural disorder encoded by overlapping genes. Genetics, 210, 303–313.

Wisotsky, S. R., Kosakovsky Pond, S. L., Shank, S. D. & Muse, S. V. 2020. Synonymous site-to-site substitution rate variation dramatically inflates false positive rates of selection analyses: ignore at your own peril. Molecular Biology and Evolution.

Yang, X., Jensen, S. I., Wulff, T., Harrison, S. J. & Long, K. S. 2016. Identification and validation of novel small proteins in Pseudomonas putida. Environmental microbiology reports, 8, 966–974.

